# Smoke exposed roots causes reduced whole-plant vascular sectoriality

**DOI:** 10.1101/2022.04.28.489912

**Authors:** Mary Benoit, Spenser Waller, Stacy L. Wilder, Michael J. Schueller, Richard A. Ferrieri

## Abstract

While photosynthates are partitioned by the relative strength of young developing leaves and roots as sinks for carbon-based resources, many plants also show a close relationship between partitioning, phyllotaxy and vascular connectivity, yielding sectorial patterns of allocation. We examined whether smoke influences phloem vascular sectoriality in a model plant, sunflower (*Helianthis annuus* L.). Using radioactive ^11^CO_2_ administered to single photosynthetically active source leaves, we examined the transport behavior and allocation patterns of ^11^C-photosynthates using gamma counting and autoradiography. Soil treated with liquid smoke caused significant reductions in phloem sectoriality involving young sink leaves and roots. The resulting increase in vascular connectivity could benefit young plant performance by allowing a more uniform allocation of nutrients and/or stress signal molecules at a critical time of their growth.

**One-Sentence Summary:** Smoke-exposed roots exhibit a significant reduction in phloem sectoriality involving carbon transport to young sink leaves and roots.

## Introduction

Globally, there has been a steady rise and increased severity in wildfires due to climate change (Abatzoglou & Williams, 2016). Despite their destructive nature, often with extensive loss to personal property and land resources, fire can play an important role to restoring land resources, especially when employed in prescribed burns (Maggard et al., 2018). Most notably, it can be an important conduit for returning carbon back into the ground, in turn restoring soil health. It can also be beneficial to removing invasive plant species that might otherwise overwhelm growth of the native population (Rundel, 1981). Of course, wildfires generate an excessive amount of smoke. Studies have shown that smoke can also influence plant re-growth immediately following a wildfire event (Keeley & Pizzorno, 1986; Baldwin et. al., 1994; Baldwin & Morse, 1994; Preston & Baldwin, 1999; Van Staden et al., 2000; Nelson et al., 2009). For example, seeds lying dormant for years in the soil can be suddenly stimulated to germinate upon exposure to smoke. Most notably are the so-called “fire chasers,” or ephemeral plants such as *Nicotiana attentuata* or wild tobacco that are known to spring up in great numbers throughout the Great Basin Desert regions of the U.S. after a wildfire event (Baldwin et al., 1994; Baldwin & Morse, 1994). In addition to being able to break seed dormancy, certain chemical constituents within smoke have been implicated with altering root growth behavior and architecture (Taylor & Van Staden, 1998; Kulkarni et al., 2006; Soós et al., 2009; Abdelgadir et al., 2012; Abdollahi, 2012; Chumpookam et al., 2012; Wang et al., 2017; Zhong et al., 2020), which in turn, can alter sink strength, or demand for photosynthates while having beneficial effects on overall plant performance. For example, increased root biomass and/or alteration of root types, have been shown to improve nutrient acquisition by better enabling plants to forage for patchy resources belowground (Giehl & von Wiren, 2014).

Higher plants possess a physiological organization that is based upon the carbon economy of their parts. Here photosynthates are partitioned across long distances according to the relative strength of the sink tissues. However, in many plant species there is also a very close relationship between partitioning, phyllotaxy and vascular connectivity giving rise to sectorial patterns of resource allocation that have evolved from selective ecological pressures over time. Such vascular restrictions in long-distance transport have been shown to exist within plants for both xylem and phloem vascular tissues (Orian et al., 2005). Sectoriality is also not something that can be defined as being black and white, but rather as shades of gray. Many dicots differ considerably in their degree of sectoriality with natural variations occurring within both herbaceous and woody species (Orians et al., 2005; Marshall, 1996). One can rationalize the existence of sectoriality in nature based on its broad ecological consequences. Consider the conditions for optimal plant performance based simply on resource availability and reaction energetics. For plant performance to remain optimized, resistance to pests, diseases or other environmental factors should be variable in time and space at scales relevant to the individual plant. Take, for example, damage caused by feeding herbivores. Evidence shows that the distribution and intensity of such damage is often patchy and varied in intensity across the different plant parts (Denno & McClure, 1983). Mounting plant defenses often relies on the production of very specialized secondary metabolites acting as chemical agents against attack. These processes can be energy demanding with repercussions to growth (Baldwin & Schmelz 1994). Logically, for a plant to mount a “full-scale” defensive response seems a waste of energy and precious resources, especially if only a portion of the plant comes under attack. Since systemic induction triggering plant defenses relies on the movement of signal molecules through the vascular system, patterns of induction will vary spatially due to vascular connectivity (Schittko & Baldwin, 2003). Hence, sectoriality from an evolutionary perspective offers a means for the plant to tailor its response to the severity of herbivore attack so as not to over-tax its growth resources. Even so, there are some who would argue there are benefits to a plant’s ability to mount a full-scale defensive response especially if attack is so severe and long-lasting that a selective tailored response can risk catastrophic unrecoverable losses to tissues resulting in death (de Bobadilla, et al., 2022). In fact, defense induction in some sectorial plant species including tomato (Orians et al., 2000) and Arabidopsis (Kiefer & Slusarenko, 2003) has been shown to be highly integrated across all tissues. Hence, loss of sectoriality in whole-plant defense induction might be beneficial to the plant’s fitness, especially when considering that regions populated by young developing seedlings after a fire event can be more susceptible to attack by pests.

Finally, one must also realize that the evolutionary rules of sectoriality are not cast in stone. It has been shown that manipulation of source-sink relations can break down such vascular barriers. For example, in soybean defoliation of one sector of a plant while depodding another allowed photosynthate to cross from source leaves in one sector to pods in another (Noodén, 1978). In the present work, we examined whether smoke could influence phloem sectoriality linked to long- distance transport of photosynthates in sunflower (*Helianthis annuus* L.) as a model sectorial dicot (Alkio et al., 2002).

## Results

Here, we filled rhizoboxes with either clean sifted topsoil, or liquid smoke treated topsoil (1:200 *dil. v/v*) to grow study plants to their V8 stage of development. Sunflower roots have a taproot from which grow several secondary and tertiary roots. Smoke treatment had no apparent effect on total number of secondary roots, nor an effect on their length, but did cause a significant increase in the tertiary root length although their number was greatly diminished relative to control plants (Figure 1).

**Figure 1.**
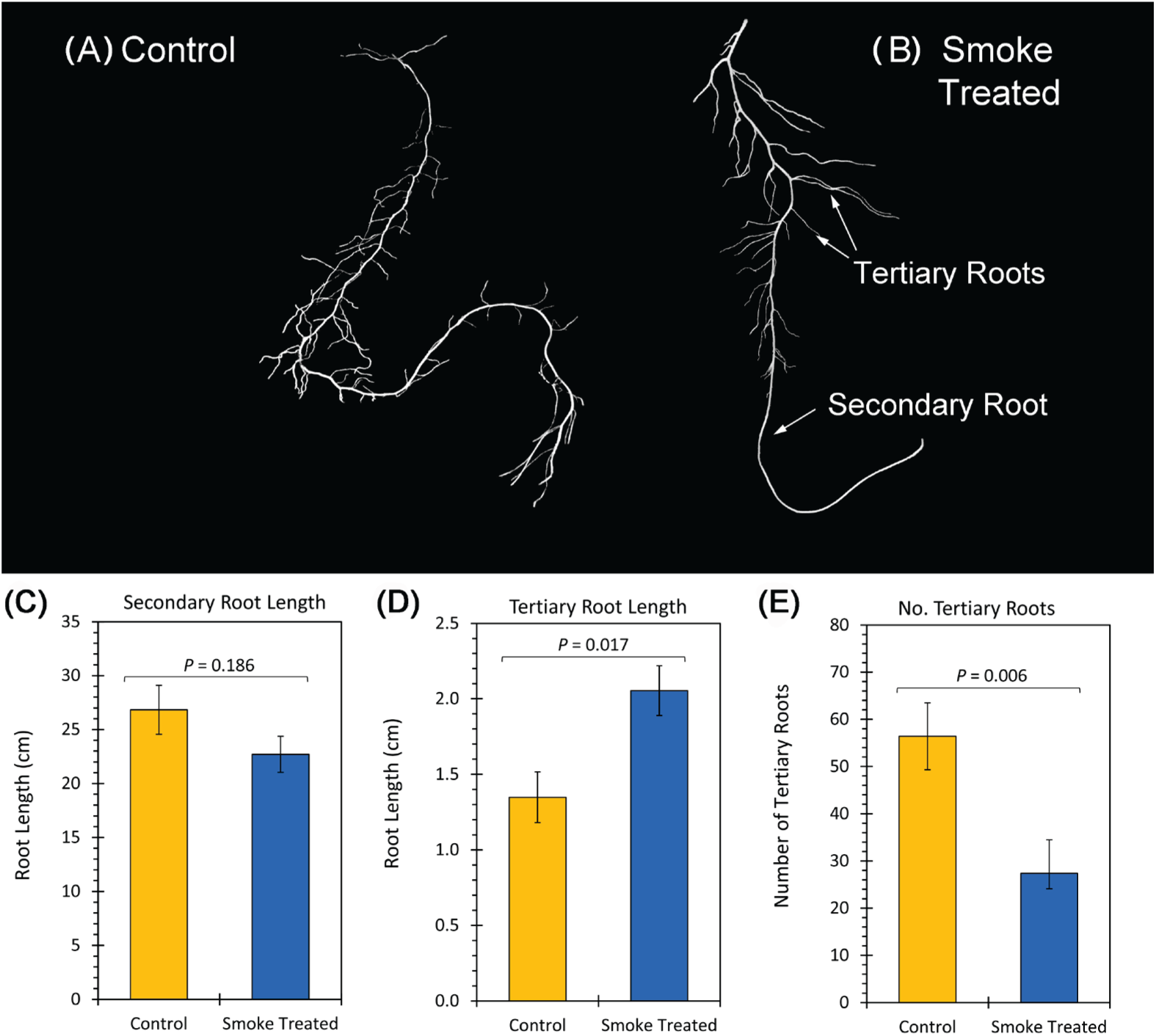
Liquid smoke treated soil at (1:200 dil. *v/v*) alters sunflower root growth traits. **(A) & (B)** Isolated secondary and tertiary roots were removed from each study plant and photographed for quantitative root tracing using AmScope software. **(C)** The length of secondary roots is presented in centimeter dimensions for control and liquid smoke treated soil growth conditions. No significant difference in root length was noted here with treatment. **(D)** The length of tertiary roots is presented in centimeter dimensions for control and liquid smoke treated soil growth conditions. Smoke treatment significantly increased tertiary root length relative to control. (**E)** The average number of tertiary roots attached to single secondary root is presented. Smoke treatment significantly reduced the number of tertiary roots attached to any single secondary root. All data represents means ± SE for N=4-6 replicates. *P*-values are shown where *P*<0.05 was statistically significant.

Radioactive ^11^CO_2_ was applied to the V4 leaf of study plants (Figure S1) where fixation and export of ^11^C-photosynthates was followed dynamically for 90 minutes. Smoke treatment did not affect ^11^CO_2_ fixation, nor did it affect leaf export or root allocation of ^11^C-photosynthates. However, the speed at which ^11^C-photosynthates moved through the taproot doubled when roots were exposed to smoke (Figure 2). Finally, autoradiography applied to V1/V2 sink leaves and to roots revealed a significant reduction in phloem sectoriality within both tissue types (Figure 3).

**Figure 2.**
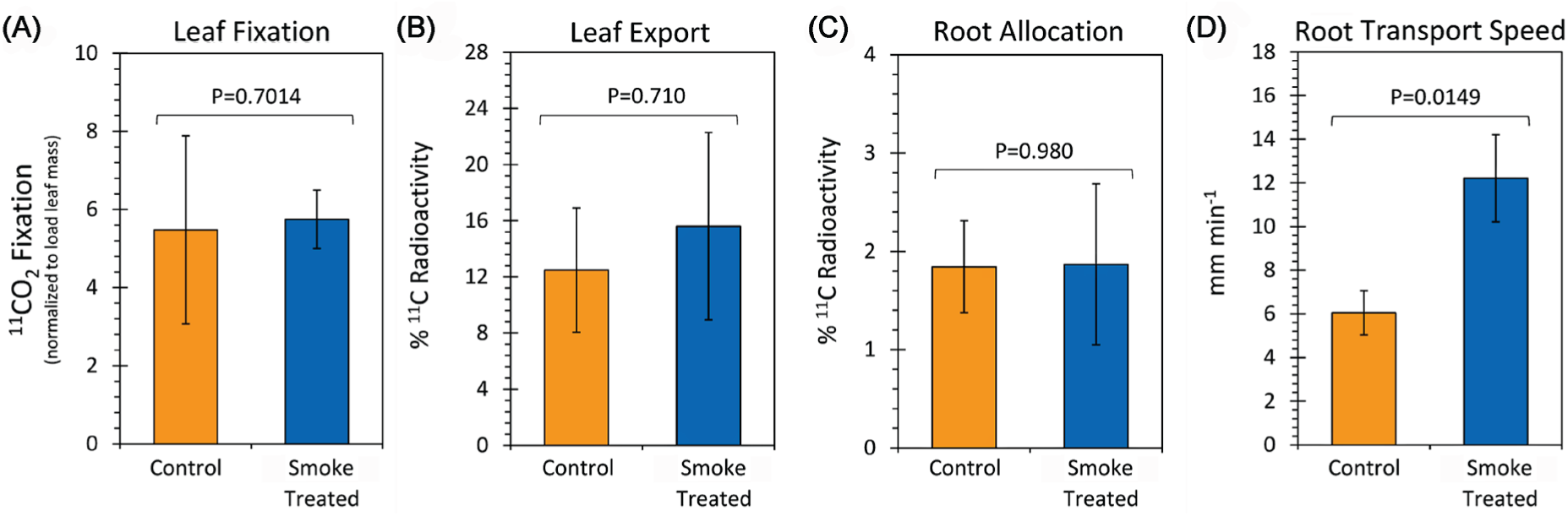
Whole-plant physiological measurements depicting carbon input, transport, and allocation. **(A)** Source leaf fixation of ^11^CO_2_ is shown as a function of soil treatment. Fixation values represent percent of the ^11^CO_2_ pulse sent to the leaf cell where data was normalized to a common leaf mass in that cell. Results indicate that liquid smoke treated soil did not affect ^11^CO_2_ fixation. **(B)** Source leaf export of ^11^C-photosynthates over a 90-minute period is presented as percent ^11^C-radioactivity fixed by the plant as a function of soil treatment. Results show that liquid smoke treatment did not affect photosynthate export from the target source leaf. **(C)** The relative percent of plant ^11^C-activity allocated to the root biomass over a 90-minute period is shown as a function of soil treatment. Results show that liquid smoke treated soil did not affect photosynthate allocation to roots as a function of soil treatment. **(D)** The transport speed of ^11^C-photosynthates within the taproot are shown in mm min-1 rate as a function of soil treatment. Results indicate that liquid smoke treated soil significantly increased the rate of transport nearly 2-fold. All data reflects mean values ± SE for N=4 replicates. *P*-values are indicated in each panel where *P*<0.05 was statistically significant.

**Figure 3.**
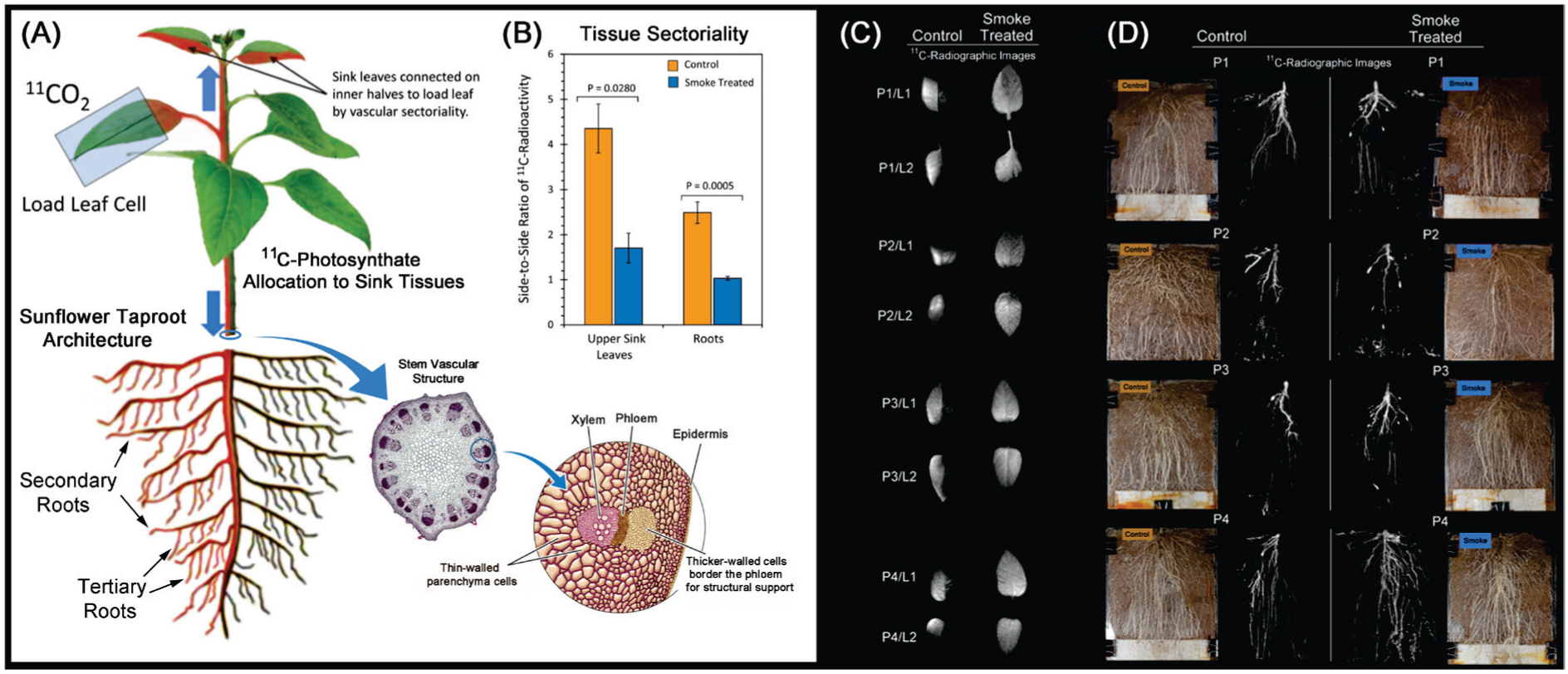
Phloem vascular connectivity of a young sunflower plant. **(A)** ^11^CO_2_ is administered to a single V4 The flow of ^11^C-photosynthates is shown in red with direct connections to only half of two younger sink leaves attached immediately above that source leaf. Flow is also shown to secondary and tertiary roots growing out from the taproot on the same side as that source leaf. The anatomical arrangement of phloem and xylem are shown as insets. **(B)** The degree of tissue sectoriality is depicted in this graph as the side-to-side ratio of ^11^C-activity measured in the two young sink leaves and in the roots. The midrib and taproot were used as the dividing lines for activity measurements using ImageQuant™ software applied to ^11^C-radiographs. A value of 1 in the graph reflects no sectoriality. Data represents means ± SE for N=4 replicates. Values of *P* < 0.05 were considered statistically significant. Both sink leaves and roots showed significant reduction of phloem sectoriality. **(C)** Radiographic images of V1 and V2 sink leaves are shown for study plants (labeled as P1-4/L1-2) subjected to control and liquid smoke (1:200 dil. *v/v*) soil treated growth conditions. **(D**) Radiographic images of roots are shown adjacent to their respective digital photographs showing the entire root mass within each rhizobox for study plants subjected to control and smoke treated soils.

## Discussion

Loss of sectoriality can have far reaching implications to plant performance especially at an early developmental stage where integrated defense induction could be critical to plant survival. As plants continue to develop, their ability to uniformly allocate carbon-based resources from source leaves to all sink tissues could aid in over-all growth performance, especially if shading becomes problematic with overlapping leaves of older plants. However, for smoke application to become a practical management practice in agriculture or forestry, more studies are needed to become a practical management practice in agriculture or forestry, more studies are needed to identify which chemical constituents in smoke are responsible for loss of sectoriality.

Aside from the possible practical applications of smoke treatment described here, this work also provides new knowledge on a more fundamental level. We note that as the “effective” root sink strength is essentially doubled due to the elimination of the sectorial vascular barriers, the speed at which carbon moves through the taproot tissue also increases proportionally. What is remarkable about this is the supply of carbon from the source leaves remains unchanged. This challenges our fundamental thinking of plant carbon allocation and source-sink relations where according to existing dogma established 90 years ago in the Münch Hypothesis (Münch, 1930), the supply of carbon in leaves is what drives a mass gradient of sugar in the phloem propelling bulk flow in long-distance transport to sink tissues. However, according to findings recently published (Babst et al., 2022) this fundamental theory is being challenged. Our findings support this challenge in that the speed of movement within the phloem appears independent of a mass gradient.

## Materials and methods

### Plant Growth

A 1 L volume of screened topsoil was treated with 200 mL of 1:200 dilution of liquid smoke (Wright’s Liquid Smoke Concentrate, B&B Foods, Inc., NJ) in deionized water then dried in an oven at 70 °C for 4 days before rescreening. Non-treated soil was washed with an equivalent amount of deionized water then dried and screened in the same manner.

Rhizoboxes were constructed from 0.0125-inch-thick plastic sheets of Plexiglass™ separated by a thin elastomer gasket made from screen spline that was super glued to the edge. Rhizobox dimensions measured 8 × 12 inches and were 0.0125-inch thick on the inside spacing after assembly. A small notch cut into the top side of the rhizobox gasket provided stem access and a ready port for administering Hoagland’s nutrient (PhytoTechnology Laboratories, Shawnee Mission, KS, USA). Rhizoboxes were filled with soil in readiness for transplanting sunflower seedlings. Sunflower seeds (Mammoth variety: Green Garden Products, Norton, MA, USA) were germinated in ProMix using 4-inch plastic pots. Five days after germination, seedlings were transplanted to the rhizoboxes. Blotter paper was placed at the bottom of each rhizobox to wick water up into the soil during plant growth. The front and rear plates were clamped tight using standard stationary binder clips and the entire assembly was wrapped in aluminum foil to prevent algae growth. Plants were placed within trays of water and placed in a commercial growth chamber (model PGC-15: Percival Scientific, Inc., Perry, IA, USA). Growth conditions for included 12- hour photoperiods, 500 μmol m^-2^ s^-1^ light intensity, and temperatures of 25°C/ 20°C (light/dark) with relative humidity at 40%. Hoagland’s nutrient (10 mL) was injected through the stem breach every 5 days.

### Production and Administration of Radioactive ^11^CO_2_

Isotope was produced on the GE 800 Series PETtrace Cyclotron located at the Missouri Research Reactor Center using high-pressure research grade N_2_ gas target irradiated with a 16.4 MeV proton beam to generate ^11^C *via* the ^14^N(p,α)^11^C nuclear transformation. The ^11^CO_2_ was trapped on the molecular sieve, desorbed, and quickly released into an air stream at 200 mL min^-1^ as a discrete pulse for labeling a V4 source leaf affixed within a 5 × 10 cm leaf cell to ensure a steady level of fixation. Plants were maintained under the same light intensity and temperature as what they experienced in the growth chamber. The load leaf affixed within the cell was pulse-fed ^11^CO_2_ for 1 min, then chased with normal air for the duration of exposure. A PIN diode radiation detector (Carroll Ramsey Associates, Berkeley, CA USA) attached to the bottom of the leaf cell enabled continuous measurement of radioactivity levels within the cell during the initial pulse and in the minutes directly following to give information on ^11^CO_2_ fixation and leaf export of ^11^C-photosynthates.

After ^11^CO_2_ pulsing, plants were incubated for 90-minutes beyond the initial tracer pulse. A continual stream of air was passed through the leaf cell during this time while levels of radioactivity were monitored using two radiation detectors (Eckler & Ziegler, Inc., Berlin, Germany 1-inch NaI gamma ray detector with photomultiplier tube) affixed 3-inches apart in lead shielding. Each detector had a field-of-view along the taproot region of the rhizobox (Figure S1). These detectors provided information on the dynamic transport as ^11^C-photosynthate moved along that portion of the root system. Data were acquired at a 1 Hz sampling rate using 0–1 V analog input into an acquisition box (SRI, Inc, Torrance, CA, USA). Measurement of ^11^C radioactivity was performed using gamma counting and data was decay-corrected to end of bombardment.

### Autoradiography

Subsequent to ^11^C dynamic measurements examining transport V1/V2 sink leaves were harvested and autoradiographs acquired by exposing phosphor plate films for approximately 10-minutes. Likewise, the stem was cut away from the rhizobox and the top plate was removed to expose the entire root mass growing on the soil surface. A phosphor plate was laid down across the entire root mass and exposed for approximately 30-minutes. Phosphor plates were read using the Typhoon 9000 imager (Typhoon™ FLA 9000, GE Healthcare, Piscataway, NJ, USA). Images were used for quantitatively determining spatial patterning of radiotracer within the two connected sink leaves and the roots. ImageQuant™ 8.2 TL software (Cytiva Life Sciences, Inc., Marlborough, MA) was used for image analysis. For sink leaves regions of interest for radioactivity measurement were manually traced in the software using the midrib as the dividing line between the left-side and right-side leaf halves. Similarly, the taproot was manually traced dividing left-side root mass from right-side root mass for radioactivity measurement.

### Tissue Gamma Counting

After imaging, whole-plant tissues were divided into three components including: (i) the V4 source leaf to which ^11^CO_2_ was administered; (ii) all other leaves and stem; and (iii) total root mass. These components were counted for ^11^C radioactivity using a NaI gamma counter. The individual components were summed together for total plant ^11^C radioactivity and corrected for radioactive decay of the isotope (t_½_ 20.4 min) back to a common zero time. Individual components were used to calculate leaf export and root allocation fractions.

### Root Phenotyping

After whole root tissue counting isolated secondary roots were removed, suspended in a tray of water, and photographed using a DSLR camera. By suspending the root in water, this allowed the finer tertiary roots to separate making it easier to document root length. Root photographs were processed using AmScope v4.11.18421 software (AmScope, Inc., Irvine, CA, USA) to determine the average length of secondary and tertiary roots, as well as the number of tertiary roots affixed to an isolated secondary root.

### Statistical Analysis

Data was analyzed using the simple Student’s t-test for comparisons of data between control and smoke treated soils (N = 4 replicates each) on each parameter measured. Statistical significance was set at P<0.05.

## Acknowledgements

The authors wish to acknowledge Pharmalogics, Inc. staff at MURR who operate the GE PETtrace cyclotron that produces carbon-11.

## Author contributions

Conceptualization – RAF; Methodology – RAF, MB; Investigation – MB, SW, MJS; Visualization - RAF, MJS; Project administration - RAF, SLW; Supervision - RAF, MJS; Writing – original draft: RAF, MJS, SLW; Writing – review & editing: MB, SW, SLW, MJS, RAF.

## Supplementary Materials

Supplemental Figure S1. Experimental setup for ^11^C tracer studies.

## Supplementary Materials for

**Fig. S1.**
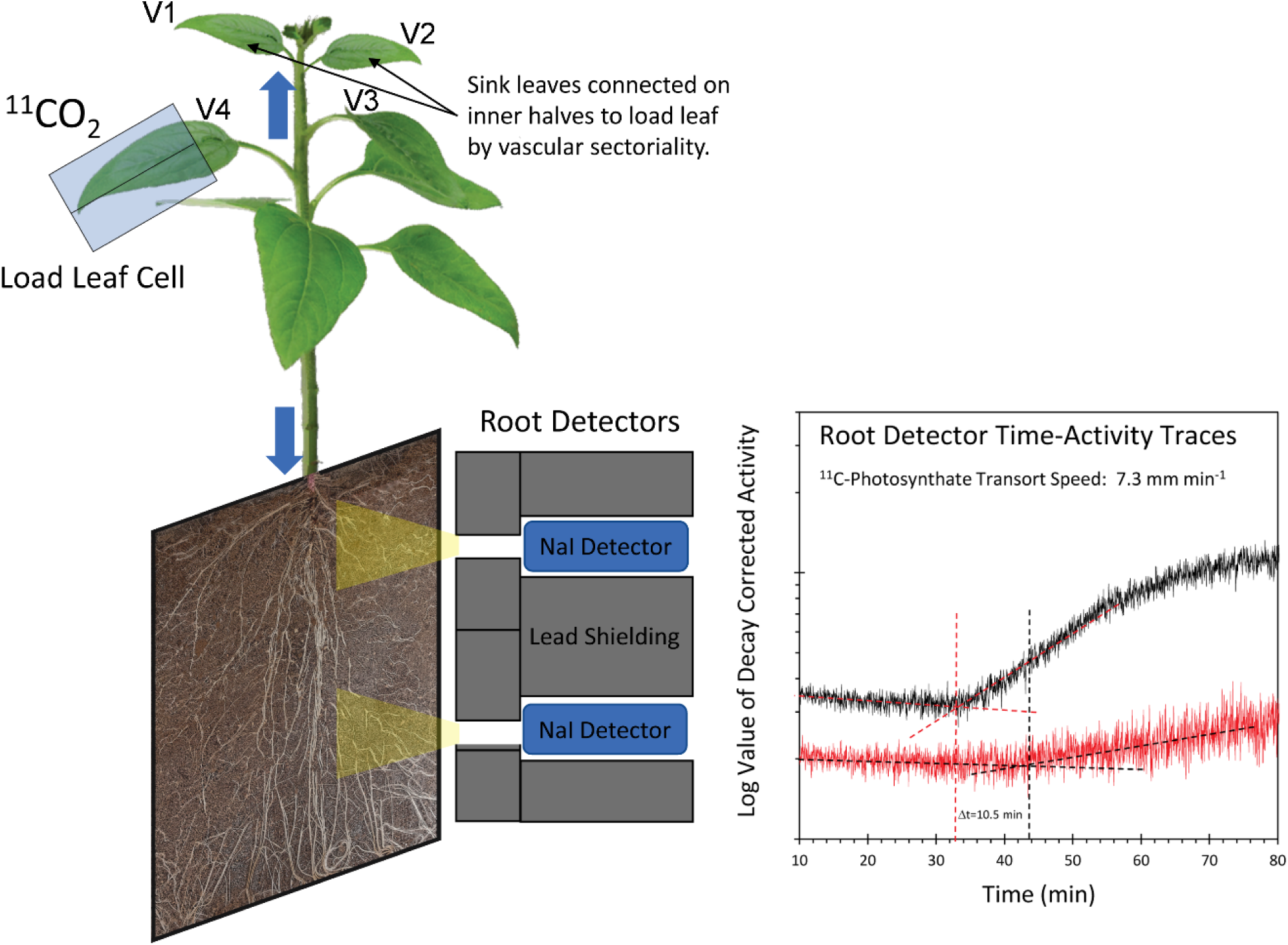
Experimental setup for ^11^C tracer studies. Source leaf V4 was captured within a gas tight leaf cell through which air was passed at 200 mL min^-1^. The ^11^CO_2_ tracer was administered as a discrete pulse within this stream of air. A PIN diode radiation detector affixed within the leaf cell provided information on fixation during the pulse. Sensitive NaI gamma radiation detectors mounted in a fixed geometry of lead shielding provided a narrow field-of-view of the taproot region for time-radioactivity measurements. Decay corrected radioactivity plotted as a function of elapsed time from pulse provided a measure of the time-to-rise which was mathematically determined by intersecting regression lines. From this data, the ^11^C-photosynthate transport speed within the taproot was calculated.

